# Identification of novel antiviral drug combinations in vitro and tracking their development

**DOI:** 10.1101/2020.09.17.299933

**Authors:** Aleksandr Ianevski, Rouan Yao, Svetlana Biza, Eva Zusinaite, Andres Männik, Gaily Kivi, Anu Planken, Kristiina Kurg, Eva-Maria Tombak, Mart Ustav, Nastassia Shtaida, Evgeny Kulesskiy, Eunji Jo, Jaewon Yang, Hilde Lysvand, Kirsti Løseth, Valentyn Oksenych, Per Arne Aas, Tanel Tenson, Astra Vitkauskiene, Marc P. Windisch, Mona Høysæter Fenstad, Svein Arne Nordbø, Mart Ustav, Magnar Bjørås, Denis Kainov

**Affiliations:** Department of Clinical and Molecular Medicine, Norwegian University of Science and Technology, 7028 Trondheim, Norway; Institute of Technology, University of Tartu, 50090 Tartu, Estonia; Icosagen Cell Factory OÜ Kambja vald Tartumaa Estonia; Institute for Molecular Medicine Finland, FIMM, University of Helsinki 00014, Finland; Applied Molecular Virology Laboratory, Institut Pasteur Korea, Sampyung-dong 696, Bundang-gu, Seongnam-si, Gyeonggi-do, Korea; Department of Laboratory Medicine, Lithuanian University of Health Science, 44307 Kaunas, Lithuania; Department of Medical Microbiology, St. Olavs Hospital, 7006 Trondheim, Norway; Department of immunology and transfusion medicine, St. Olavs Hospital, 7006 Trondheim, Norway

**Author notes:** (A.I.); (R.Y.); (S.B.); (H.L.); (K.L.); (V.O.); (P.A.A.); (M.B.); (S.A.N.). (E.Z.); (T.T.); (N.S.). (M.U.Jr.); (M.U.); (A.M.); (G.K.); (A.P.); (K.K.); (E.M.). (E.K.). (M.W.), (E.J.); (J.Y.). (A.V.). (M.H.F.).

**Keywords:** antivirals, antiviral drug combinations, broad-spectrum antivirals, virus

## Abstract

Combination therapies have become a standard for the treatment for HIV and HCV infections. They are advantageous over monotherapies due to better efficacy and reduced toxicity, as well as the ability to prevent the development of resistant viral strains and to treat viral co-infections. Here, we identify several new synergistic combinations against emerging and re-emerging viral infections *in vitro*. We observed synergistic activity of nelfinavir with investigational drug EIDD-2801 and convalescent serum against SARS-CoV-2 infection in human lung epithelial Calu-3 cells. We also demonstrated synergistic activity of vemurafenib combination with emetine, homoharringtonine, gemcitabine, or obatoclax against echovirus 1 infection in human lung epithelial A549 cells. We also found that combinations of sofosbuvir with brequinar and niclosamide were synergistic against HCV infection in hepatocyte derived Huh-7.5 cells, whereas combinations of monensin with lamivudine and tenofovir were synergistic against HIV-1 infection in human cervical TZM-bl cells. Finally, we present an online resource that summarizes novel and known antiviral drug combinations and their developmental status. Overall, the development of combinational therapies could have a global impact improving the preparedness and protection of the general population from emerging and re-emerging viral threats.

## 1. Introduction

Every year, emerging and re-emerging viruses, such as S-ARS-CoV-2, MERS-CoV, Zika virus (ZIKV), Ebola virus (EBOV), influenza A virus (FLUAV), and Rift Valley fever virus (RVFV) surface from natural reservoirs to infect, disable, and kill people [1, 2]. As of September 2020, the number of people infected with SARS-CoV-2 continues to rise, with a global death toll of around 1 million. Developing novel vaccines and antiviral drugs against emerging and re-emerging viruses is time-consuming and costly, deterring many drug developers and researchers from pursuing them until it is too late.

Drug repurposing, also called drug repositioning, is a strategy for generating additional value from an existing drug by targeting diseases other than that for which it was originally intended [37,38]. This strategy has significant advantages over new drug discoveries, since chemical synthesis steps, manufacturing processes, reliable safety and pharmacokinetic profiles in preclinical and early clinical developmental phases (phase 0, I and IIa) have already been completed. Therefore, drug repositioning for treatment COVID-19 provides unique translational opportunities, including a substantially higher probability of reaching the market as compared with developing new virus-specific drugs and vaccines, as well as significantly reduced costs and timelines to clinical availability [30,39,40].

Many viruses, however, easily develop resistance to single drug use [3]. Combination therapies can lower the evolution of drug-resistant viral variants by attacking the virus using multiple mechanisms. Moreover, combinations may lead to increased antiviral activity in a synergistic or additive manner, leading to lower drug dosage requirements to achieve the same effect. These low doses reduce the toxicity of each drug in combination and their adverse reactions. Additionally, combinations of two or more antivirals could be administered in a cocktail to target a broad range of viruses. Such combinations could serve as front line therapeutic option against poorly characterized emerging viruses, re-emerging drug-resistant viral strains or viral co-infections. For example, a cocktail of broad-spectrum nelfinavir and remdesivir could potentially be developed for the treatment of at least 18 human viruses (drugvirus.info). Thus, antiviral drug combinations may become a standard treatment of emerging and re-emerging viral infections.

Indeed, antiviral drug combinations have become a standard treatment of rapidly evolving viruses, such as HIV and HCV (https://www.drugs.com/drug-class/antiviral-combinations.html). These include abacavir/dolutegravir/lamivudine (Triumeq), darunavir/cobicistat/emtricitabine/tenofovir (Symtuza), ledipasvir/sofosbuvir, sofosbuvir/velpatasvir, and lopinavir/ritonavir (Kaletra). Of note, Kaletra has also been tested against SARS-CoV-2 infections in clinical studies (ChiCTR2000029308). Many other drug combinations are now in clinical trials against SARS-CoV-2, HCV, HBV, HSV-1, and other viral infections (NCT04291729, ChiCTR2000030894, NCT03111108, NCT01045278, NCT00255034, NCT02480166, NCT00383864, NCT01023217, NCT00922207, NCT02360592, NCT00922207, etc.).

Our recent *in vitro* studies have revealed antiviral synergism across a wide range of compounds and viruses. For example, we have found that a combination of pimodivir with gemcitabine was synergistic against FLUAV infection in human macrophages [4]. We also demonstrated that combinations of obatoclax and saliphenylhalamide at nanomolar concentrations have a synergistic effect against ZIKV infection in retinal pigment epithelium (RPE) cells [5]. In addition, we showed that a combination of nelfinavir with salinomycin, amodiaquine, obatoclax, emetine, or homoharringtonine exhibits synergistic antiviral activity against SARS-CoV-2 in kidney epithelial cells extracted from an African green monkey (Vero-E6) [6].

Here, we report that combinations of nelfinavir with investigational drug EIDD-2801 and convalescent serum were synergistic against SARS-CoV-2 infection in human lung epithelial Calu-3 cells. We further showed that combinations of vemurafenib, together with emetine, homoharringtonine, gemcitabine, and obatoclax were synergistic against echovirus 1 infection in human lung epithelial A549 cells. Combinations of sofosbuvir with brequinar and niclosamide were also shown to be effective against HCV infection in hepatocyte-derived Huh-7.5 cells. Finally, combinations of monensin with lamivudine and tenofovir were shown to have antiviral synergy against HIV-1 infection in human cervical TZM-bl cells. To summarize the scientific and clinical information regarding available and emerging antiviral drug combinations, we also present a freely accessible web resource that we have built, which aggregates all the currently known synergistic interactions between antiviral agents.

## 2. Materials and Methods

### 2.1. Drugs

Supplementary Table S1 lists compounds, their suppliers, and catalogue numbers. To obtain 10 mM stock solutions, compounds were dissolved in dimethyl sulfoxide (DMSO; Sigma-Aldrich, Germany) or milli-Q water. Collection of convalescent serum from patients recovered from COVID-19 (SARS-CoV-2_2) has been described in our previous study [6]. The solutions were stored at -80 °C until use.

### 2.2. Cell Cultures

Human non-small-cell lung cancer Calu-3 and RPE cells were grown in DMEM-F12 supplemented with 10% FBS, 100 μg/mL streptomycin, and 100 U/mL penicillin (Pen-Strep). Human adenocarcinoma alveolar basal epithelial A549 and Vero-E6 cells were grown in DMEM supplemented with 10% FBS and Pen-Strep. The cell lines were maintained at 37 °C with 5% CO_2_. ACH-2 cells, which possess a single integrated copy of the provirus HIV-1 strain LAI (NIH AIDS Reagent Program), were grown in RPMI-1640 medium supplemented with 10% FBS and Pen/Strep. TZM-bl, previously designated JC53-bl (clone 13) is a human cervical cancer HeLa cell line, stably expressing the firefly luciferase under control of the HIV-1 LTR promoter. TZM-bl cells were grown in DMEM supplemented with 10% FBS and Pen/Strep. The human hepatoma Huh-7.5 cell line was grown in DMEM supplemented with 10% FBS, non-essential amino acids (NEAA), L-glutamine and Pen/Strep [7]. All cell lines were grown in a humidified incubator at 37 °C in the presence of 5% CO_2_.

### 2.3. Viruses

The SARS-CoV-2 hCoV-19/Norway/Trondheim-S15/2020 strain has been described in our previous study [6]. It was amplified in a monolayer of Vero-E6 cells in the DMEM media containing Pen/Strep and 0.2% bovine serum albumin. EV1 (Farouk strain; ATCC) was provided by Prof. Marjomäki from University of Jyväskylä [19]. EV6 was from Kainovs’ laboratory [8]. The viruses were amplified in a monolayer of A549 cells in the DMEM media containing Pen/Strep and 0.2% BSA. For the production of HIV-1, 6 × 10^6^ ACH-2 cells were seeded in 10 mL medium. Virus production was induced by the addition of 100 nM phorbol-12-myristate-13-acetate. The cells were incubated for 48 h. The HIV-1-containing medium was collected. The amount of HIV-1 was estimated by measuring the concentration of HIV-1 p24 in the medium using anti-p24-ELISA, which was developed in-house. Recombinant purified p24 protein was used as a reference. Cell culture-derived infectious HCV (HCVcc) was produced as described before [9]. Briefly, Huh-7.5 cells transiently transfected with HCV RNA transcripts of a cell culture-adapted JFH1 genome expressing NS5A-GFP fusion protein (JFH1_5 A/5B_GFP). HCVcc containing medium was collected at 4 days post-transfection. Viral supernatant was clarified by filtration using a syringe filter with a 0.2 μm pore size (Millipore, Bedford, MA). All virus stocks were stored at −80 °C.

### 2.4. Neutralization Assay

Approximately 4 × 10^4^ Vero-E6 cells were seeded per well in 96-well plates. The cells were grown for 24 h in DMEM supplemented with 10% FBS and Pen-Strep. Serum sample was prepared in 3-fold dilutions at 7 different concentrations, starting from 40 μg/mL in the virus growth medium (VGM) containing 0.2% BSA and Pen-Strep in DMEM. Virus hCoV-19/Norway/Trondheim-S15/2020 was added to achieve a multiplicity of infection (moi) of 0.1 and incubated for 1h at 37 °C. 0.1 % DMSO was added to the control wells. The Vero-E6 cells were overplayed with VGM containing mixture of the virus and convalescent serum. After 72 h the medium was removed, and a CellTiter-Glo assay (Promega, Sweden) was performed to measure cell viability.

### 2.5. Drug Test

Approximately 4 × 10^4^ Vero-E6, A549, or RPE cells were seeded per well in 96-well plates. The cells were grown for 24 h in DMEM or DMEM-F12 supplemented with 10% FBS and Pen-Strep. The medium was replaced with DMEM or DMEM-F12 containing 0.2% BSA and Pen-Strep. The compounds were added to the cells in 3-fold dilutions at 7 different concentrations, starting from 30 μM. No compounds were added to the control wells. The cells were mock- or virus-infected at a moi of 0.1. After 24 (EV1) or 72 (SARS-CoV-2) h of infection, the medium was removed from the cells, and a CellTiter-Glo assay was performed to measure cell viability.

The half-maximal cytotoxic concentration (CC_50_) for each compound was calculated based on viability/death curves obtained on mock-infected cells after nonlinear regression analysis with a variable slope using GraphPad Prism software version 7.0a (GraphPad Software, San Diego, CA, USA). The half-maximal effective concentrations (EC_50_) were calculated based on the analysis of the viability of infected cells by fitting drug dose–response curves using the four-parameter (4PL) logistic function *f*(*x*):

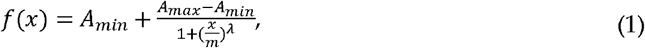

where *f*(*x*) is a response value at dose *x, A*_*min*_ and *A*_*max*_ are the upper and lower asymptotes (minimal and maximal drug effects), *m* is the dose that produces the half-maximal effect (EC_50_ or CC_50_), and *λ* is the steepness (slope) of the curve. The relative effectiveness of the drug was defined as the selectivity index (SI = CC_50_/EC_50_).

### 2.6. Virus Quantification

For testing the production of infectious virions, we titered the viruses as described in our previous studies [10-12]. In summary, media from the viral culture were serially diluted from 10^−2^ to 10^−7^ in serum-free media containing 0.2% bovine serum albumin (BSA). The dilutions were applied to a monolayer of Vero-E6 (for SARS-CoV-2) or A549 (for EV1) cells in 24-well plates. After one hour, cells were overlaid with virus growth medium containing 1% carboxymethyl cellulose and incubated for 72 (for SARS-CoV-2) or 48 h (for EV1). The cells were fixed and stained with crystal violet dye, and the plaques were calculated in each well and expressed as plaque-forming units per mL (pfu/mL).

### 2.7. Drug Combination Test and Synergy Calculations

Vero-E6 cells were treated with different concentrations of two drugs and infected with SARS-CoV-2 (moi 0.1) or mock. After 72 h, cell viability was measured using CellTiter-Glo. A549 cells were treated with different concentrations of two drugs and infected with EV1 (moi 0.1) or mock. After 24 h, cell viability was measured using CellTiter-Glo. TZM-bl cells were treated with different concentrations of two drugs and infected with HIV-1 (corresponding to 300 ng/mL of HIV-1 p24) or mock. After 48 h post infection (hpi), the media was removed from the cells, the cells were lysed, and firefly luciferase activity was measured using the Luciferase Assay System (Promega, Madison, WI, USA). In a parallel experiment, cell viability was measured using CellTiter-Glo. We also examined toxicity and antiviral activity of drug combinations using GFP-expressing HCV in Huh-7.5 cells by following previously described procedures [9].

To test whether the drug combinations act synergistically, the observed responses were compared with expected combination responses. The expected responses were calculated based on the ZIP reference model using SynergyFinder version 2 [13, 14]. We quantified synergy scores, which represent the average excess response due to drug interactions (i.e., 10% of cell survival beyond the expected additivity between single drugs has a synergy score of 5).

### 2.8. Gene Expression Analysis

A549 cells were treated with 10 μM vemurafenib or vehicle at indicated concentrations. Cells were infected with EV1 at moi 0.1 or mock. After 8 h, total RNA was isolated using RNeasy Plus Mini kit (Qiagen, Hilden, Germany). Gene expression profiling was done using Illumina Human HT-12 v4 Expression BeadChip Kit according to manufacturer’s recommendation as described previously [4]. Raw microarray data were normalized using the BeadArray, and Limma packages from Bioconductor suite for R. Normalized data were further processed using variance and intensity filter. Genes differentially expressed between samples and controls were determined using the Limma package. Benjamini-Hocberg multiple correction testing method was used to filter out differentially expressed genes based on a q-value threshold (q < 0.05). Filtered data were sorted by logarithmic fold change (log2FC). Heatmap was generated using an in-house developed interface, Breeze [15]. Gene set enrichment analysis was performed using open-source software (www.broadinstitute.org/gsea)

### 2.9. Cytokine Profiling

The medium from EV1- or mock-infected, non- or drug-treated A549-cells was collected at 24 hpi and clarified by centrifugation for 5 min at 14,000 rpm. Cytokines were analyzed using Proteome Profiler Human Cytokine Array Kit (R&D Systems) according to manufacturer’s instructions.

### 2.10. Metabolic Analysis

Metabolomics analysis was performed as described previously [4]. Briefly, 10 μL of labeled internal standard mixture was added to 100 μL of the sample (cell culture media). Next, 0.4 mL of solvent (99% ACN and 1% FA) was added to each sample. The insoluble fraction was removed by centrifugation (14,000 rpm, 15 min, 4°C). The extracts were dispensed in OstroTM 96-well plate (Waters Corporation, Milford, USA) and filtered by applying vacuum at a delta pressure of 300–400 mbar for 2.5 min on Hamilton StarLine robot’s vacuum station. The clean extract was collected to a 96-well collection plate, placed under the OstroTM plate. The collection plate was sealed and centrifuged for 15 min, 4000 rpm, 4° C, and placed in auto-sampler of the liquid chromatography system for the injection. Sample analysis was performed on an Acquity UPLC-MS/MS system (Waters Corporation, Milford, MA, USA). The autosampler was used to perform partial loop with needle overfill injections for the samples and standards. The detection system, a Xevo TQ-S tandem triple quadrupole mass spectrometer (Waters, Milford, MA, USA), was operated in both positive and negative polarities with a polarity switching time of 20 ms. Electrospray ionization (ESI) was chosen as the ionization mode with a capillary voltage at 0.6 KV in both polarities. The source temperature and desolvation temperature of 120° and 650°C, respectively, were maintained constant throughout the experiment. Declustering potential (DP) and collision energy (CE) were optimized for each compound. Multiple Reaction Monitoring (MRM) acquisition mode was selected for quantification of metabolites with individual span time of 0.1 s given in their individual MRM channels. The dwell time was calculated automatically by the software based on the region of the retention time window, number of MRM functions and also depending on the number of data points required to form the peak. MassLynx 4.1 software was used for data acquisition, data handling, and instrument control. Data processing was done using TargetLynx software, and metabolites were quantified by calculating curve area ratio using labeled internal standards (IS) (area of metabolites/area of IS) and external calibration curves. Metabolomics data were log2 transformed for linear modeling and empirical-Bayes-moderated t-tests using the LIMMA package (https://bioconductor.org/packages/release/bioc/html/limma.html). To analyse the differences in metabolites levels, a linear model was fit for each metabolite. The Benjamini-Hochberg method was used to correct for multiple testing. The significant metabolites were determined at a Benjamini-Hochberg false discovery rate (FDR) controlled at 10%. The heatmap was generated using the pheatmap package (https://cran.rproject.org/web/packages/pheatmap/index.html) based on log-transformed profiling data. MataboAnalyst 3.0 was used to identify pathways related to EV1 infection (www.msea.ca). In this pathway analysis tool, the pathway data are derived from KEGG database (www.genome.jp/kegg/)

### 2.11. Website Development

The current landscape of the available and emerging antiviral drug combinations were reviewed and summarized in a database that can be freely accessed at https://antiviralcombi.info. The information for the database was obtained from PubMed, clinicaltrials.gov, DrugBank, DrugCentral, the Chinese Clinical Trials Register (ChiCTR), and EU Clinical Trials Register databases [25–27], as well as other public sources. To extend the coverage of our database, we manually review each article to extract literature-supported raw drug combination data where available. The website was developed with PHP v7 technology using D3.js v5 (https://d3js.org/) for visualization.

## 3. Results

### 3.1. Novel Anti-SARS-CoV-2 Combinations

As of September 2020, the number of people infected with severe acute respiratory coronavirus 2 (SARS-CoV-2) continues to skyrocket, with around 30 million cases worldwide. The World Health Organization (WHO) has highlighted the need for better control of SARS-CoV-2 infections. We have recently shown that serum samples from patients recovered from SARS-CoV-2 infections could neutralize the virus and prevent virus-mediated cell death [6]. Here, we tested convalescent serum sample from G614 recovered patient in combination with orally available nelfinavir, which was shown to inhibit the protease of different beta-coronaviruses, including SARS-CoV-2 [16]. We observed that nelfinavir and G614 serum displayed antiviral synergism without detectable toxicity in human lung epithelial Calu-3 cells (synergy score: 24; Fig. 1A).

**Figure 1.**
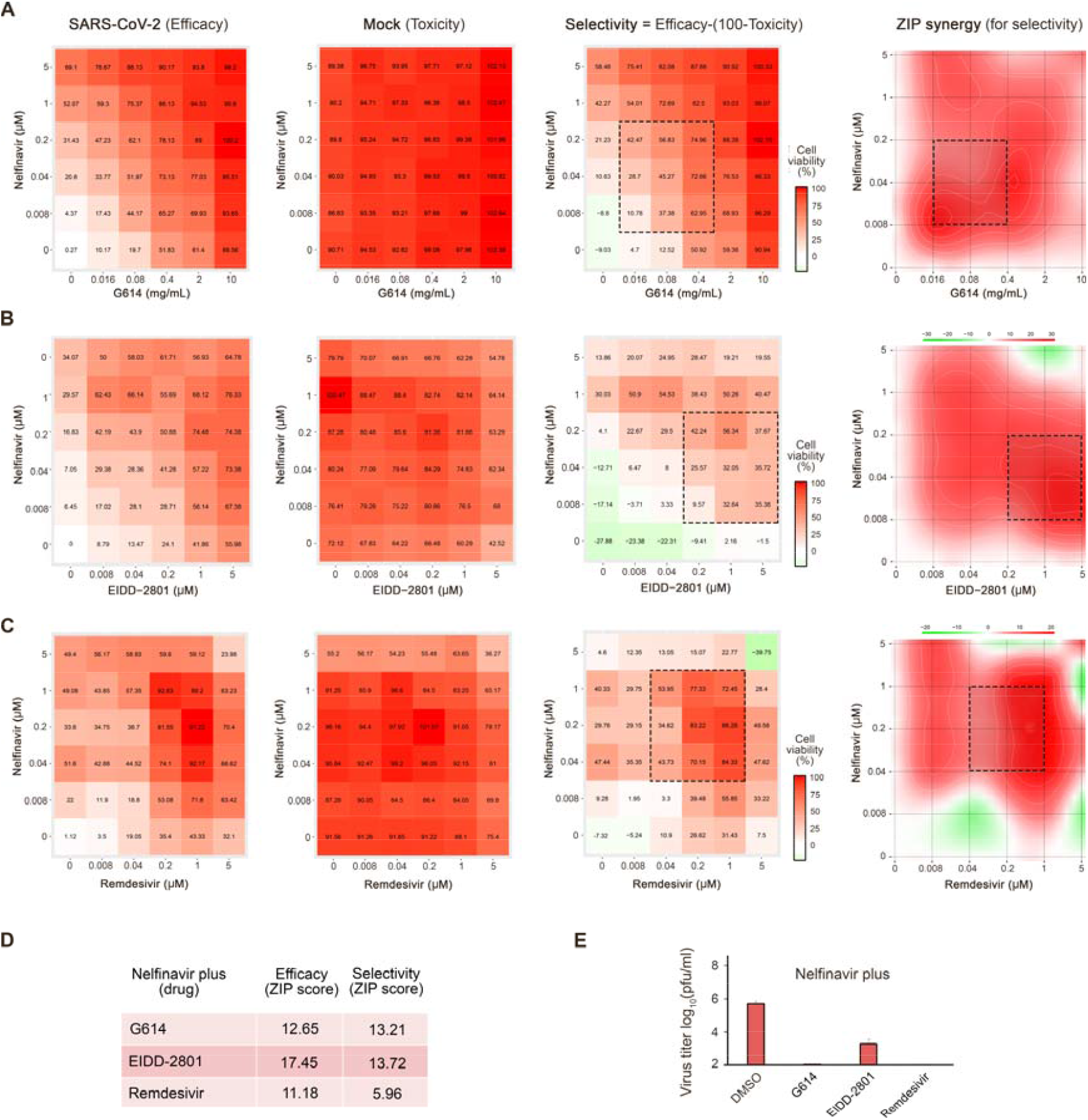
Combinations of nelfinavir with neutralizing antibody, EIDD-2801, or remdesivir rescue Calu-3 cells from SARS-CoV-2-mediated death and inhibit virus replication. (A-C) The interaction drug combination landscapes measured as 6×6 dose-response matrices using a CTG assay on SARS-CoV-2- and mock-infected cells (two left panels). The interaction landscape shows selectivity (efficacy-toxicity) and synergy for the drug combinations (right panels). (D) ZIP synergy scores calculated for efficacy and selectivity dose-response matrices for 3 drug combinations. (E) The effects of 1 μM nelfinavir plus 0,1% DMSO, 2 mg/mL G614, 1 μM EIDD-2801, or 1 μM remdesivir on viral replication measured by plaque reduction assay.

Cycloheximide, cepharanthine, EIDD-2801, and remdesivir have been reported recently to possess anti-SARS-CoV-2 activity [17-19]. We confirmed their anti-SARS-CoV-2 activities on Vero-E6 cells (Fig. S1). Moreover, a combination of nelfinavir with remdesivir has been shown recently to possess anti-SARS-CoV-2 activity in Vero-E6 cells (doi.org/10.1101/2020.06.16.153403). We tested combinations of cycloheximide, cepharanthine, and EIDD-2801 or remdesivir with nelfinavir in Calu-3 cells. We observed that only nelfinavir plus EIDD-2801 or remdesivir were synergistic at non-cytotoxic concentrations (synergy scores: 14 and 6; Fig. 1B-D). Moreover, at selected concentrations, nelfinavir combinations with convalescent serum G614, EIDD-2801, or remdesivir reduced the SARS-CoV-2 production by >2 logs in comparison to nelfinavir alone (Fig. 1E). Thus, the nelfinavir-convalescent serum G614, nelfinavir-EIDD-2801, and nelfinavir-remdesivir combinations could result in better efficacy and decreased toxicity for the treatment of SARS-CoV-2 than drugs alone.

### 3.2. Novel Anti-EV1 Combinations

EV1 belongs to the genus *Enteroviruses*. Enteroviral infections affect humans worldwide by causing the common cold, hand-foot-and-mouth disease, meningitis, myocarditis, pancreatitis, and poliomyelitis. Enteroviruses are also associated with chronic diseases such as type I diabetes, asthma, and allergies. Enteroviruses are non-enveloped viruses that belong to the family of *Picornaviridae*. They include 12 species, enterovirus A-H, and J and rhinovirus A-C.

There are no approved therapies to treat enterovirus infections. In order to identify antiviral drug candidates, we screened the FIMM oncology drug collection (527 drugs) against EV1 in human cancer lung epithelial A549 and retinal pigment epithelial RPE cells using cell viability assay as readout. We identified vemurafenib as an inhibitor of EV1 replication. Vemurafenib (marketed as Zelboraf) is an FDA-approved inhibitor of the cellular B-Raf enzyme for the treatment of late-stage melanoma. Vemurafenib interrupts the B-Raf/MEK/ERK pathway, only if the B-Raf has the common V600E mutation [20]. A549 and RPE are non-BRAF mutated cells. Importantly, vemurafenib inhibited EV1, but not EV6 virus (structural identity – 88%, similarity - 93%) in A549 cells, indicating that it targets the viral proteins (Fig. S2A).

Next, we studied the effect of vemurafenib on the metabolism of EV1- and mock-infected A549 cells. We analyzed 111 polar metabolites in cell culture supernatants at 24 hpi. We quantified 90 metabolites. EV1 infection affected the levels of adenine, adenosine, hypoxanthine, glutathione, NAD, AMP, guanosine, and sucrose. Vemurafenib had some effect on the levels of these metabolites in both EV1- and mock-infected A549 cells (Fig. S2B).

We also evaluated the effect of vemurafenib on transcription in EV-1 and mock-infected A549 cells. Cells were treated with 10 μM vemurafenib or DMSO and infected with EV1 or mock. After 8 hours, we analyzed the expression of cellular genes using RNA microarray. We found that vemurafenib deregulated transcription of several genes in EV1- and mock-infected cells (Fig. S2C). Interestingly, gene set enrichment analysis (GSEA) revealed that vemurafenib affected transcription of 17 cellular genes belonging to GO_RESPONSE_TO_OXYGEN_CONTAINING_COMPOMPOUND and GO_RESPONSE_TO_ENDOGENOUS_STIMULUS gene sets (p-value 9.95 e-15 and 1.24 e-14; FDR q-value 9.45 e-11 and 9.45 e-11). These results indicate that vemurafenib could also target cellular factor(s) involved in the transcription of antiviral genes.

In addition, we evaluated the effect of vemurafenib on the production of cytokines and growth factors in EV1-infected and non-infected cells. After 24 h the medium from EV1- or mock-infected, DMSO- or drug-treated A549 cells was collected and clarified by centrifugation. Cytokines were analyzed using Proteome Profiler Human Cytokine Array Kit. Fig. S2d shows that EV1 replication suppressed secretion of CXCL-1, PDF-AA, CCL2, IL8, Angiogenin, IGFBP-2, and VEGF and activated production of FGF-2, whereas vemurafenib treatment reversed this virus-mediated effect.

We examined whether combinations of vemurafenib with safe-in-man emetine, homoharringtonine, obatoclax, gemcitabine, or dalbavancin can protect cells from EV1-mediated death better than vemurafenib alone [21]. Virus- and mock-infected A549 cells were treated with an increasing concentration of vemurafenib and increasing concentrations of two drugs. After 24 h, cell viability was measured. We calculated the synergy scores as well as selectivity for each drug combinations considering their toxicity. Fig. 4 shows that vemurafenib plus emetine, homoharringtonine, gemcitabine, or obatoclax were synergistic (synergy score > 5). However, at selected concentrations, only two combinations (vemurafenib plus emetine or homoharringtonine) reduced the EV1 production by >2 logs in comparison to vemurafenib alone (Fig. 2C). We concluded that the addition of emetine or homoharringtonine could decrease the effective dose of vemurafenib against EV1 *in vitro*.

**Figure 2.**
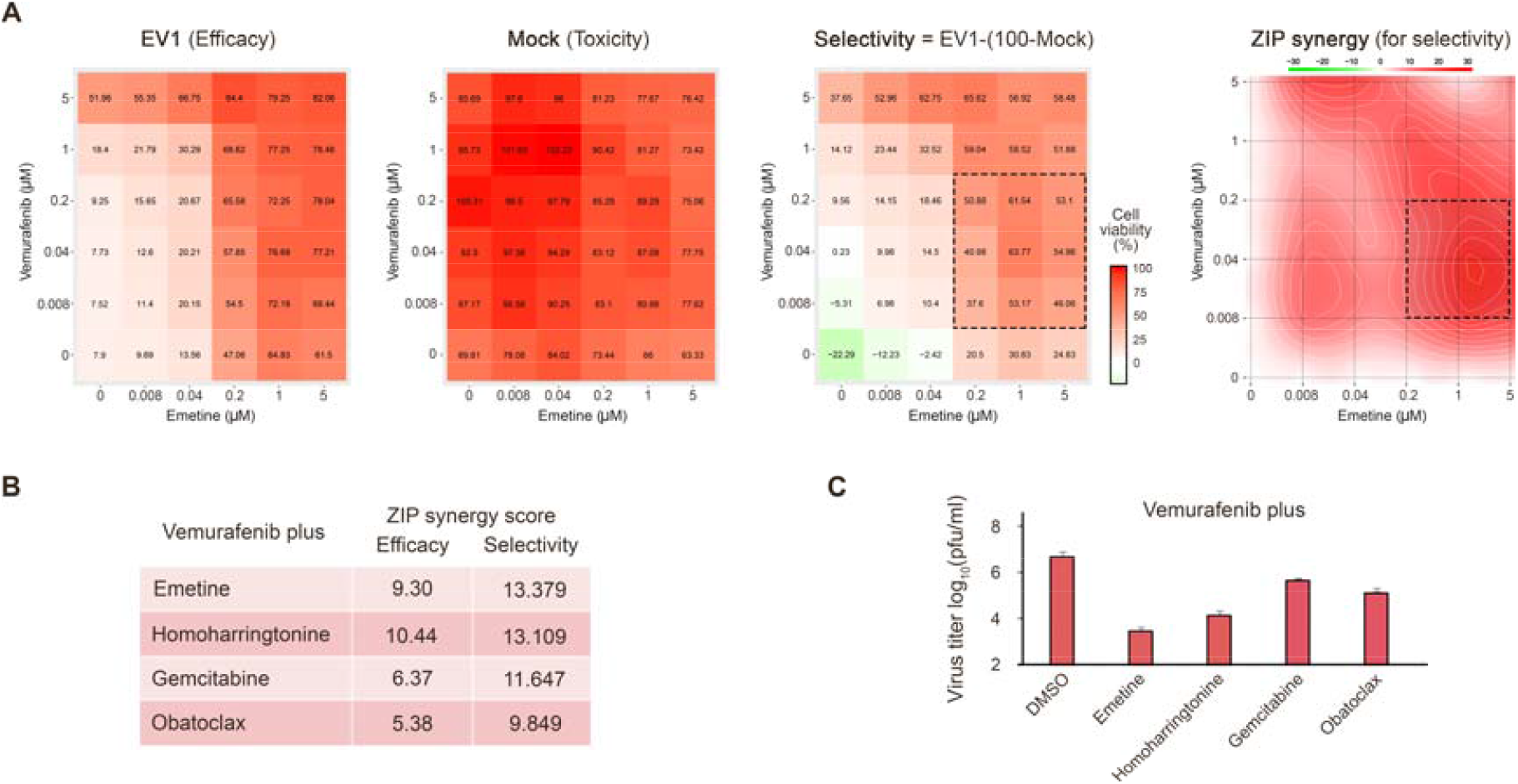
Combinations of vemurafenib with other antiviral agents rescue A549 cells from EV1-mediated death and inhibit virus replication. (A) The representative interaction landscapes of one of the drug combinations measured using a CTG assay on EV1- and mock-infected cells (left). The representative interaction landscape shows selectivity and synergy of the drug combination (right). (B) Synergy scores were calculated for efficacy (EV1-infected) and selectivity (EV1-infected - Mock) dose-response matrices for 4 drug combinations. (C) The effects of 5 μM vemurafenib plus 0,1% DMSO, 0.2 μM emetine, 0.2 μM homoharringtonine, 0.2 μM obatoclax, or 1 μL gemcitabine on viral replication measured by plaque reduction assay.

**Figure 3.**
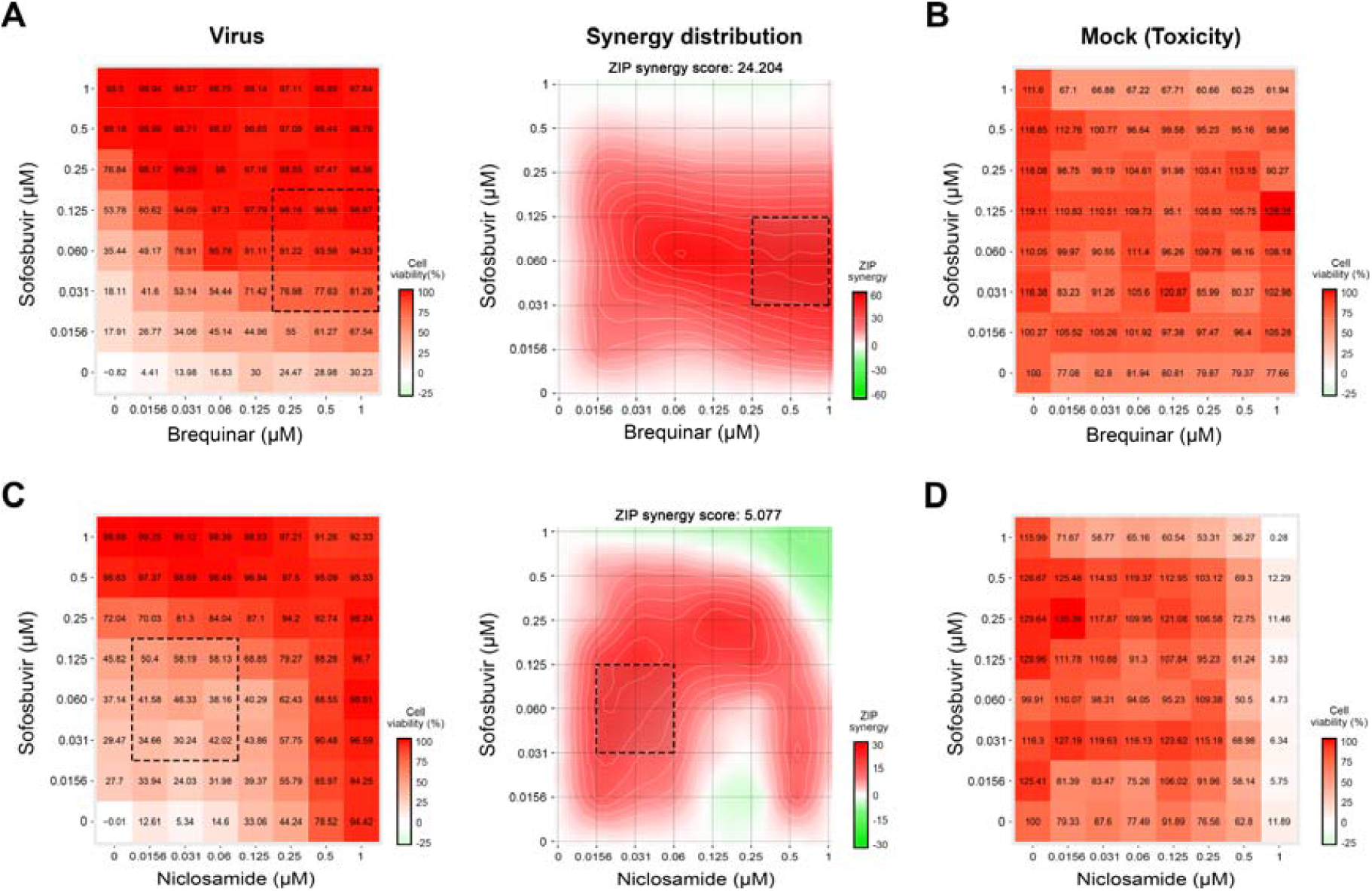
Combinations of sofosbuvir with other antiviral agents inhibit HCV infection in Huh-7.5 cells. The interaction drug combination landscapes measured as 8×8 dose-response matrices using a reporter virus expression (GFP) and CTG assay on HCV- and mock-infected cells, respectively.

**Figure 4.**
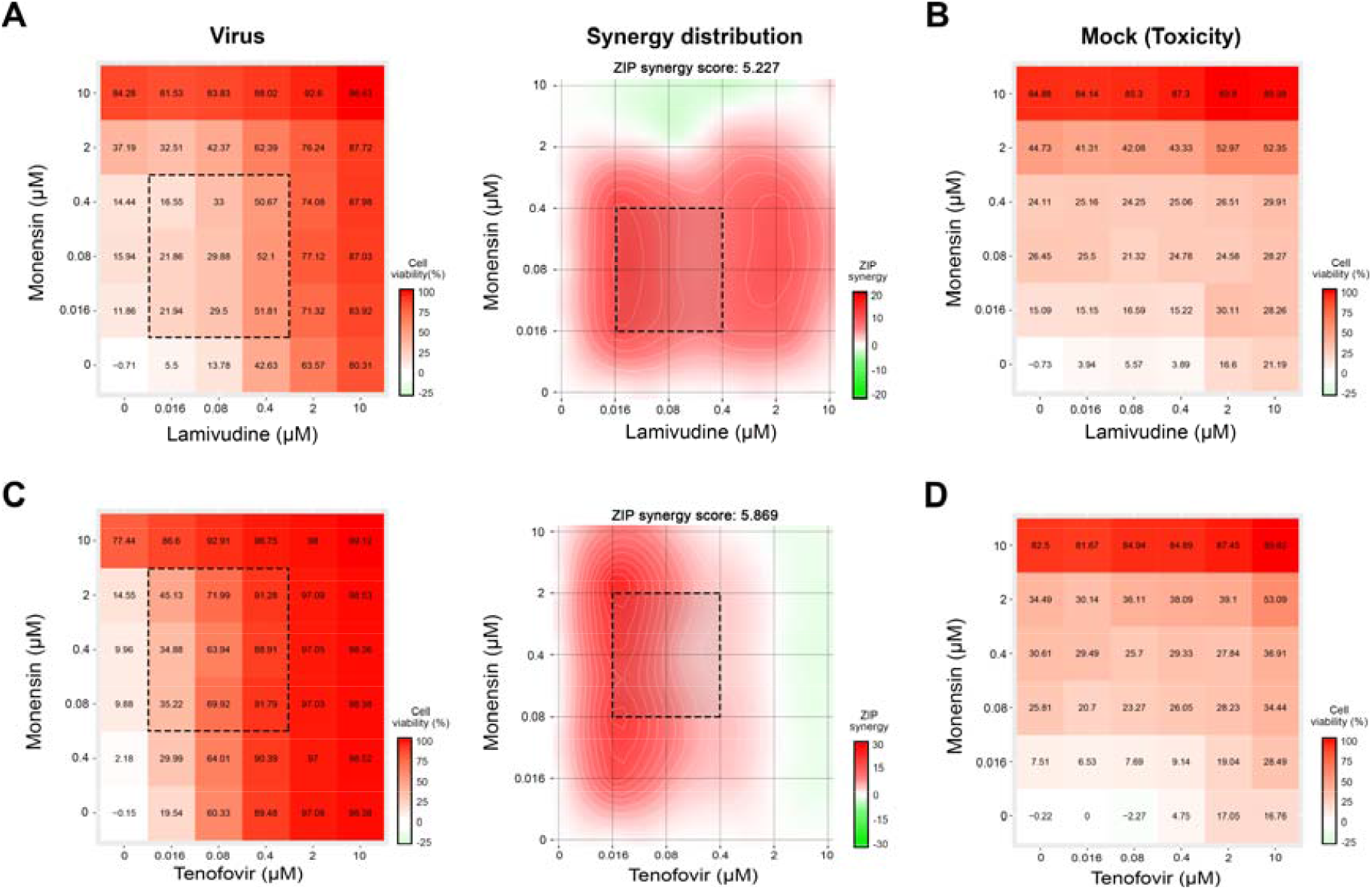
Combinations of monensin with lamivudine or tenofovir inhibit HIV-1 infection in TZM-bl cells. The interaction drug combination landscapes measured as 6×6 dose-response matrices using a reporter cell expression (luciferase) and CTG assay on HIV-1- and mock-infected cells, respectively.

### 3.3. Novel Anti-HCV Combinations

Through literature review, we identified several drugs that could be combined to inhibit HCV infection *in vitro [22-24]*. We tested combinations of sofosbuvir with brequinar, emetine, homoharringtonine, or niclosamide using GFP-expressing HCV in infected Huh-7.5 cells [23]. Eight different concentrations of the compounds alone or in combinations were added to virus- or mock-infected cells. HCV-mediated GFP expression and cell viability were measured after 48 h post-infection to determine compound efficiency and toxicity. We identified two drug combinations, which lowered GFP-expression without detectable cytotoxicity at indicated concentrations: sofosbuvir-brequinar and sofosbuvir-niclosamide, with synergy scores of 24 and 5, respectively (Fig. 3).

### 3.4. Novel drug combinations against HIV-1 infections

Through literature review, we identified several drugs that could be combined to inhibit HIV-1 infection *in vitro* [21, 22, 24]. We tested combinations of lamivudine with brequinar, suramin, ezetimibe, minocycline, rapamycin, or monensin, as well as combinations of tenofovir with the same drugs against HIV-1-mediated firefly luciferase expression in TZM-bl cells. The firefly luciferase open reading frame is integrated into the genome of TZM-bl cells under the HIV-1 LTR promoter. Six different concentrations of the compounds alone or in combinations were added to virus- or mock-infected cells. HIV-induced luciferase expression and cell viability were measured after 48 h to determine compound efficiency and toxicity. Our screen identified two combinations (lamivudine-monensin and tenofovir-monensin) that suppressed HIV-1-mediated firefly luciferase expression without detectable cytotoxicity with synergy scores of 5.2 and 5.9, respectively (Fig. 4).

### 3.5. Drug combination database

To rapidly respond to emerging and re-emerging viral diseases, we developed a freely accessible database summarizing antiviral drug combinations and their developmental status. The website is updated regularly and incorporates novel combinations as they emerge or change the statuses of existing ones as updates occur. The database comprises 985 antiviral drug combinations (Fig. 5A), including 2- and 3-drug cocktails. It covers 612 unique drugs and 68 different viruses (Fig. 5B).

**Figure 5.**
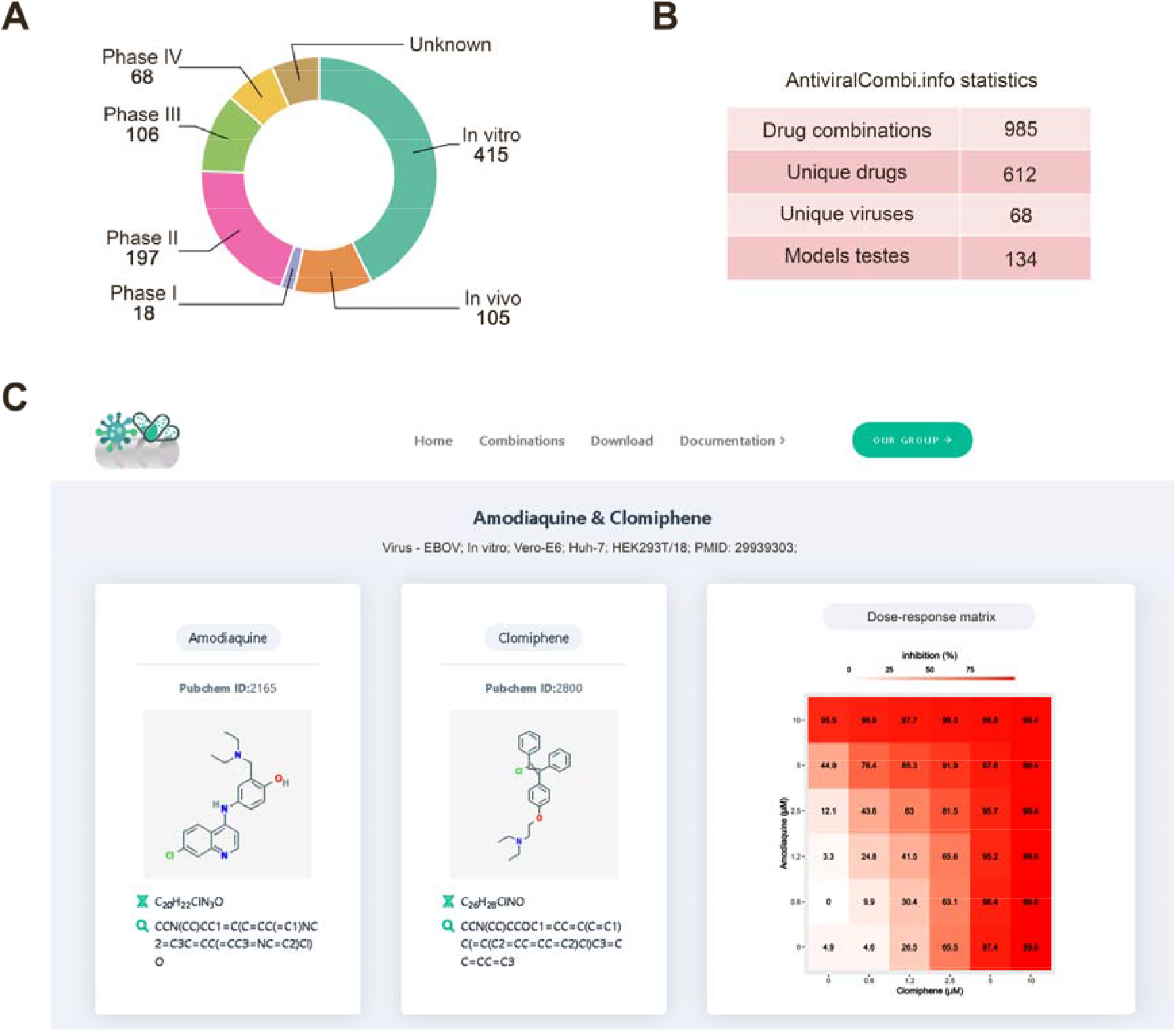
AntiviralCombi.info database summarizing existing antiviral drug combinations. A. Developmental phases of combinations are presented in the database. B. Database summary statistics. C. An example snapshot of the database showing information on the amodiaquine and clomiphene drug combination.

To extend the coverage of our database, we manually review each article to extract literature-supported raw drug combination data that were available and calculated the most common synergy scores (ZIP, Bliss, Loewe, and HSA) for those. This allows filtering drug-pairs to select the most synergistic combinations for selected viruses and interactively investigate their raw combination data (Fig. 5C). The website also includes various other functional modules, including ‘search,’ ‘filtering,’ ‘graphical visualization,’ and ‘download’ modules.

## 4. Discussion

Drug combinations are emerging as useful tools to treat infections because of their increased efficacy, decreased toxicity, and the ability to prevent the development of antiviral-resistant strains of viruses. Because drug combinations can be used to boost the effectiveness of existing antiviral drugs, they represent an attractive and innovative way to find effective treatments for viral diseases that do not currently have effective antiviral options. Here, we have reported several novel combinations that show synergism and have better efficacy than single drug therapies.

Our previous experiments have uncovered synergism between nelfinavir and several other drugs against SARS-CoV-2 in Vero-E6 cells [25]. Nelfinavir is an orally active drug that was approved for use in the treatment of HIV infection, meaning that its safety profile in humans is already understood. Nelfinavir was also shown to target the SARS-CoV-2 protease indicating that it can quickly be moved to clinical trials to treat COVID-19 (doi.org/10.26434/chemrxiv.12332678.v2). In this paper, we report novel combinations with nelfinavir that could have a substantial effect against SARS-CoV-2 infection. In particular, we have used convalescent serum and nelfinavir to achieve high synergism. This is particularly important because the FDA has granted emergency use authorization for convalescent serum therapy for the treatment of COVID-19, and a recent safety study has found that its use is generally safe in patients [26]. Therefore, due to its promising antiviral activity and its ability to be rapidly adopted as a therapy, we believe that further research on combinations of convalescent plasma or neutralizing antibodies such as BD-368-2 (doi.org/10.1016/j.cell.2020.09.035) with nelfinavir or other viral inhibitors is warranted.

We have also uncovered synergism between nelfinavir and EIDD-2801. EIDD-2801 is an orally administered investigational antiviral that was originally developed for the treatment of influenza, while remdesivir is an intravenously administered, approved broad-spectrum antiviral agent that has already been widely used to treat patients with COVID-19 [27, 28]. EIDD-2801 is a nucleotide analog. Because EIDD-2801 has already been under development for the treatment of SARS-CoV-2 infection and are well-positioned to respond to the COVID-19 pandemic, we strongly urge further investigation into these combinations to further improve the antiviral efficacy of these drugs.

We also identified four novel synergistic antiviral interactions which successfully inhibited EV1 infection. EV1 belongs to the *Enterovirus* genus, which currently has no approved antiviral therapies and can cause a range of illnesses, including the common cold, hand foot and mouse disease, meningitis, myocarditis, pancreatitis, and poliomyelitis. Of the four combinations we uncovered, vemurafenib-ementine and vemurafenib-homoharringtonine stood out by inhibiting EV1 replication by over 2 orders of magnitude more than inhibition by vemurafenib alone.

Vemurafenib is an approved, orally available drug that was originally developed for the treatment of melanoma. Recently, it was also shown to have some antiviral activity against influenza A virus infection [29]. Our studies identified and characterized the antiviral activity of vemurafenib against EV1 and suggested that the drug may also have broad-spectrum antiviral properties by targeting virus and host factors (www.drugvirus.info). We also showed that the antiviral effect of vemurafenib is amplified by combination with emetine and homoharringtonine. These compounds have been used to treat human disease and are therefore understood to be safe for human use. Thus, we recommend further development of these two drug combinations as novel anti-EV1 therapeutics.

Combination therapy is already widely used in the treatment of HCV and HIV. Our studies have identified four novel synergistic drug combinations against HCV and HIV. These combinations boost the activity of existing approved antivirals by adding other broad-spectrum antiviral agents that were not originally indicated for these purposes. In the case of HCV, we combined sofosbuvir (an FDA-approved anti-HCV drug) with brequinar (an investigational anti-cancer agent) and niclosamide (an approved antihelminthic agent). In the case of HIV, we combined monensin (a veterinary antibiotic) with lamivudine and tenofovir (both approved anti-HIV agents). Consequently, we have uncovered several ways to improve the efficacy of standard therapies by taking advantage of the time- and cost-saving benefits of drug repositioning (Fig. 6).

**Figure 6.**
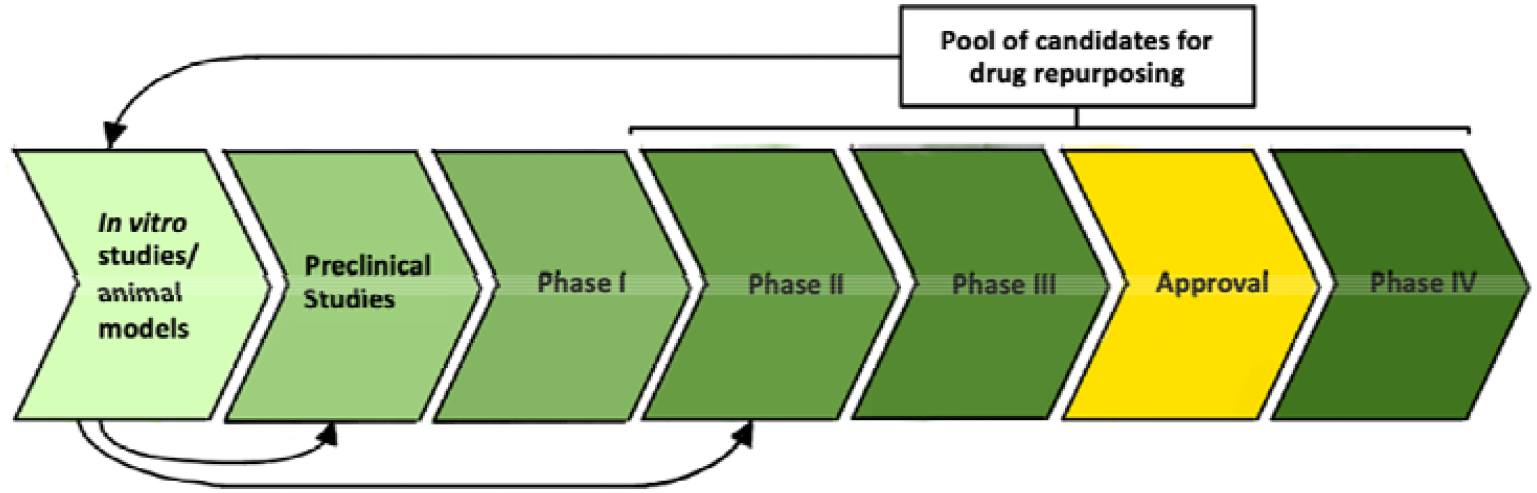
Repurposing safe-in-man antiviral combinations for treatment of viral infections. Repositioned drugs often come from the pool of approved and investigative drugs that have already passed phase I clinical trials. *In vitro* studies can uncover new antiviral activity of these drugs, either through drug synergism with other drugs or by monotherapy. Once proof-of-concept has been established *in vitro* and *in vivo*, repositioned drugs may be able to skip preclinical PK/PD and toxicology testing and phase I clinical trials, depending on the existing body of knowledge around the drug. Thus, repurposed drugs may easier proceed to phase II clinical trials, allowing for a cheaper and faster path to market.

Although the use of drug cocktails is not new, many patients and doctors still rely on monotherapies to fight viral infections [30]. Much of this is due to incomplete knowledge of drug interactions. Thus far, research on drug synergy is often disjointed, and there has yet to be a centralized system for aggregating and summarizing synergistic, additive, and antagonistic interactions between antiviral drugs.

To underscore the potential benefits of synergistic drug combinations and provide an organized summary of all currently known synergistic antiviral drug combinations, we constructed a freely accessible web resource that can be accessed at https://antiviralcombi.info. We hope that by aggregating synergism data uncovered by our group and other researchers and drug developers can be easily directed to investigate the most promising combinations to develop the most effective antiviral treatment strategies possible.

## 5. Conclusions

In this paper, we have identified novel and reviewed known synergistic antiviral drug combinations in a publicly accessible database. In particular, our study has uncovered several combinations that could be used to treat SARS-CoV-2, EV1, HIV-1, and HIV infections. Our goal is to complete preclinical studies and translate our findings into trials in patients. The most effective and tolerable combinations will have a global impact, improving the protection of the general population from emerging and re-emerging viral infections or co-infections and allowing the swift management of drug-resistant strains. Our bigger ambition is to find potential biomarkers for the prediction of novel drug combinations for the treatment of emerging and re-emerging viral infections. This can be used as means for the fast and cheaper identification of safe and effective antiviral options.

## Supplementary Materials

The following information is available online: **Table S1:** Compounds, their suppliers, and catalogue numbers. **Figure S1**. Recombinant antibodies from patients recovered from COVID-19 as well as remdesivir, EIDD-2801, cycloheximide, and cepharanthine prevent the virus-mediated death of Vero-E6 cells. **Figure S2**. Vemurafenib is a novel inhibitor of EV1 replication.

## Author Contributions

All authors contributed to the methodology, software, validation, formal analysis, investigation, resources, data curation, writing, and review and editing of the manuscript. D.K. conceptualized, supervised, and administrated the study and acquired funding. All authors have read and agreed to the published version of the manuscript.

## Acknowledgments

We thank Dr. Vidya Velagapudi for metabolomics analysis, Dr. Gaute Brede for useful discussions, and personal of Biomedicum function genomics (FuGen) unit for transcriptomics analysis.

## Supplementary information

**Table S1.**
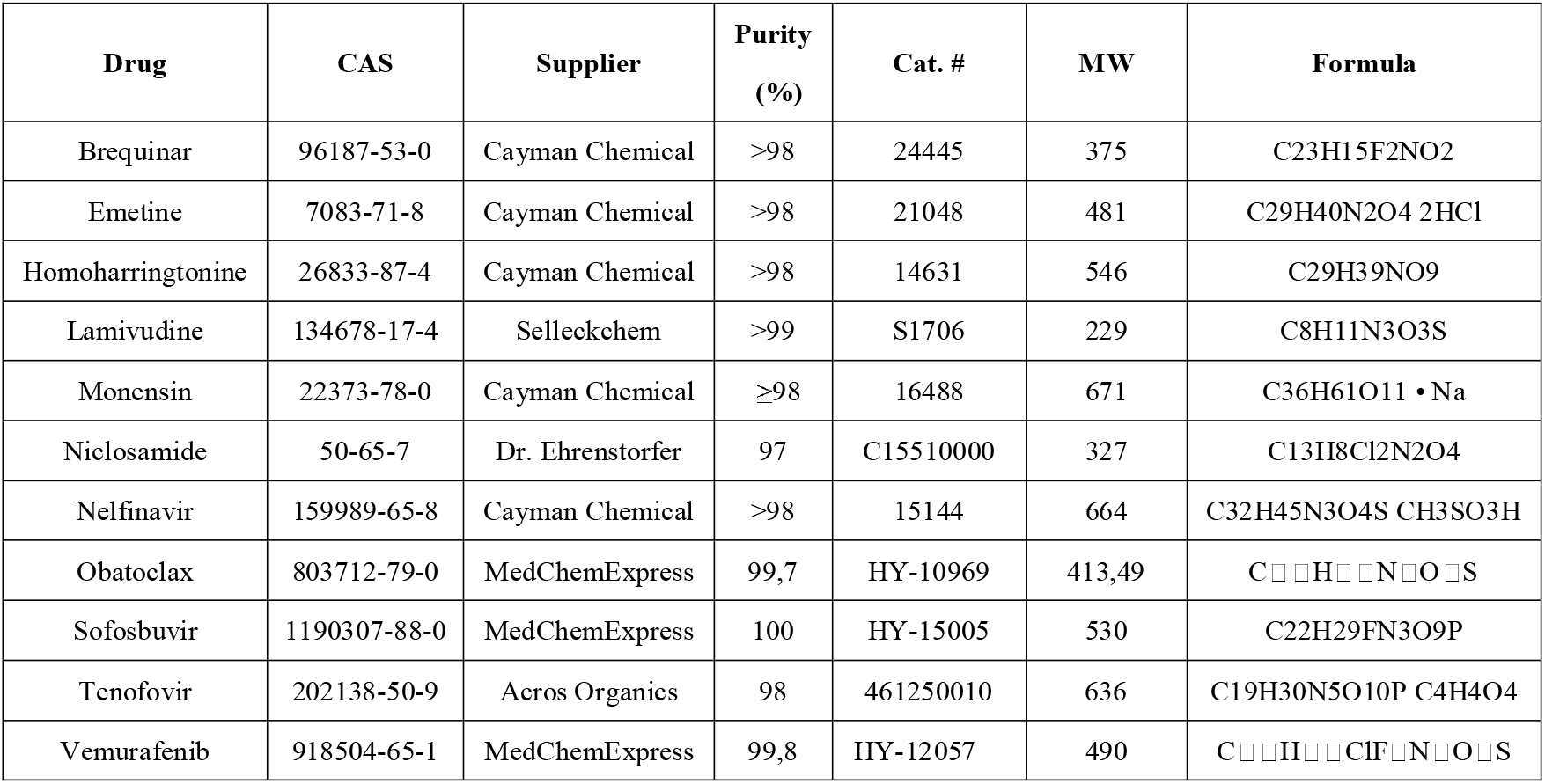
Compounds, their suppliers, and catalogue numbers.

**Fig. S1.**
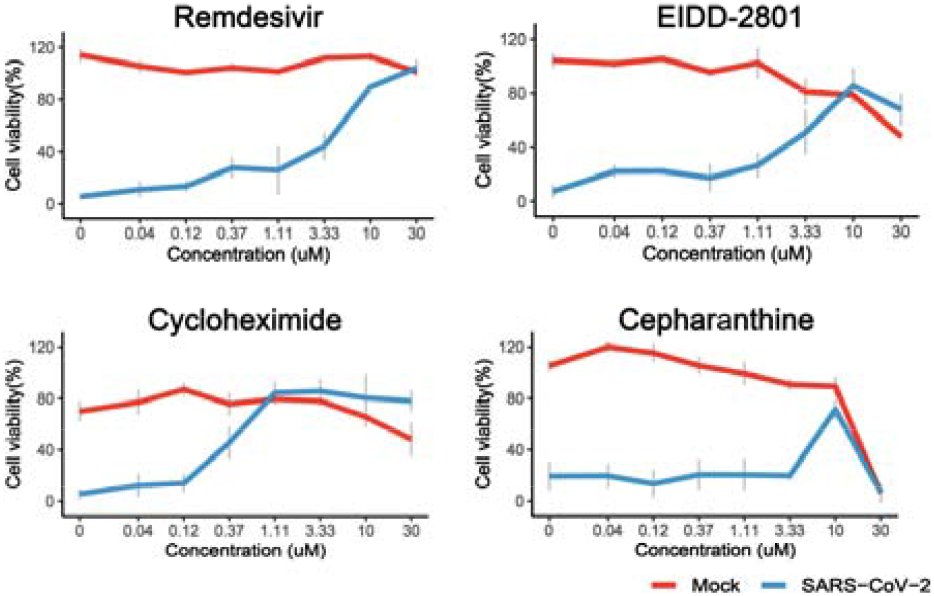
Remdesivir, EIDD-2801, cycloheximide, and cepharanthine prevent the virus-mediated death of Vero-E6 cells. The cells were treated with increasing concentrations of a compound and infected with the HCoV-19/Norway/Trondheim-S15/2020 strain (moi, 0.1) or mock. After 72 h, cell viability was determined using CellTiter-Glo. Mean ± SD; n = 3.

**Fig. S2.**
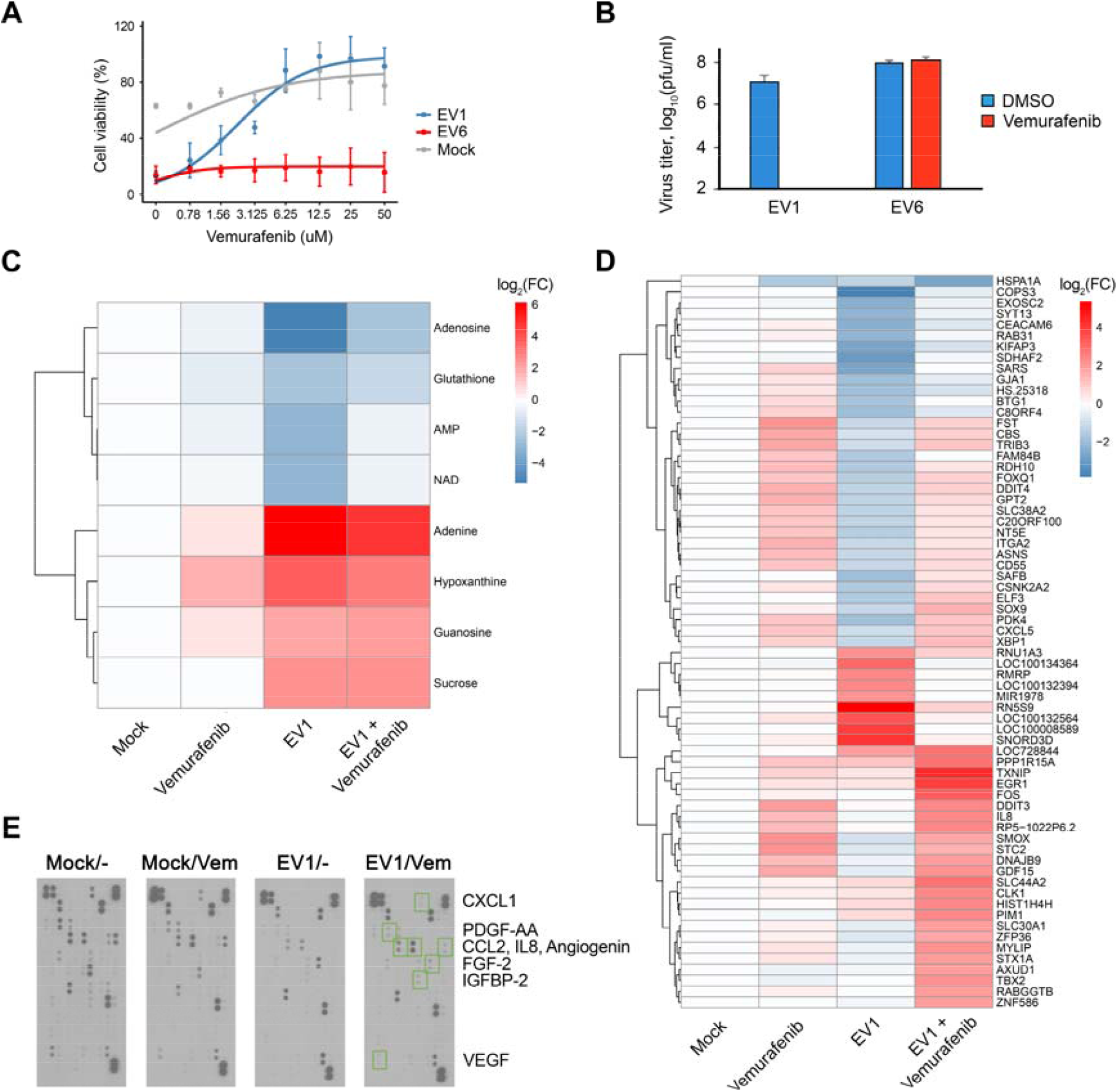
Vemurafenib is a novel inhibitor of EV1 replication. (A) A549 cells were treated with increasing concentrations of vemurafenib or DMSO and infected with EV1 (moi, 0.1), EV6 (moi, 0.1) or mock. The viability of the cells was determined at 24 hpi with the Cell Titer Glo assay (mean ± SD; n = 3). (B) EV1 and EV6 produced in A549 cells treated with vemurafenib (10 μM) or DMSO was tittered using plaque reduction assay, and the viral titers were calculated (mean ± SD; n = 3). (C) A549 cells were treated with 10 μM vemurafenib or non-treated, infected with EV1 (moi 0.1) or mock. After 24 h, cell culture supernatants were collected, and levels of 111 polar metabolites were determined using mass spectrometry. A heatmap of the most variable metabolites is shown (cut-off: log2FC > 1.5 and <−1.5). Rows represent metabolites, columns represent samples. Each cell is colored according to the log2-transformed values of samples, expressed as fold-change (FC) relative to the average of mock controls. Metabolites are ranked based on the FC (mean; n = 3). (D) A549 cells were treated with 10 μM vemurafenib or solvent and infected with EV1 (moi 0.1) or mock. At 8 hpi, the cells were collected, total RNA was extracted, and gene expression profiling was carried out. A heatmap of the most variable genes is shown (cut-off: log2FC > 1 and <−1). Rows represent gene symbols, columns represent treatments. Each cell is coloured according to the log2-transformed and quantile-normalized values of the samples, expressed as FC relative to the average of mock controls. (E) A549 cells were treated with 10 μM vemurafenib or non-treated, infected with EV1 (moi 0.1), or mock. After 24 h, cell culture media were collected, and relative levels of cytokines, chemokines, growth factors, and other soluble proteins were determined using a human cytokine array. Scanned array membranes are shown, and dots corresponding to affected cytokines are indicated.

